# Fine-tuning the transcriptional regulatory model of adaptation response to phosphate stress in maize (*Zea mays* L.)

**DOI:** 10.1101/2021.01.18.427100

**Authors:** Pranjal Yadava, Vikram Dayaman, Astha Agarwal, Krishan Kumar, Ishwar Singh, Rachana Verma, Tanushri Kaul

## Abstract

The post green revolution agriculture is based on generous application of fertilizers and high-yielding genotypes that are suited for such high input regimes. Cereals, like maize (*Zea mays* L.) are capable of utilizing less than 20% of the applied inorganic phosphate (Pi) - a non-renewable fertilizer resource. A greater understanding of the molecular mechanisms underlying the acquisition, transportation and utilization of Pi may lead to strategies to enhance phosphorus use efficiency (PUE) in field crops. In this study, we selected 12 Pi responsive genes in maize and carried out their comparative transcriptional expression in root and leaf tissues of a hydroponically grown Pi stress tolerant maize inbred line HKI-163, under sufficient and deficient Pi conditions. Pi starvation led to significant increase in root length; marked proliferation of root hairs and lesser number of crown roots. Eleven genes were significantly up or down regulated in Pi deficient condition. The putative acid phosphatase, *ZmACP5*, expression was up regulated by 162.81 and 74.40 fold in root and leaf tissues, respectively. The RNase, *ZmRNS1* showed 115 fold up regulation in roots under Pi deprivation. Among the two putative high affinity Pi transporters *ZmPht1;4* was found specific to root, whereas *ZmPht2* was found to be up regulated in both root and leaf tissues. The genes involved in Pi homeostasis pathway (*ZmSIZ1, SPX1* and *Pho2*) were up regulated in root and leaf. In light of the expression profiling of selected regulatory genes, an updated model of transcriptional regulation under Pi starvation in maize has been proposed.

## Introduction

Phosphorus (P) is one of the most important macronutrient for plant growth and development (Raghothama 1999, Bindraban et al 2020). It is required for the constitution of many cellular components, including nucleic acids, membranes, etc. and participates in enzymatic reactions and signal transduction processes. In the soil, P exists in inorganic, organic and phytate forms. Plants acquire P by their roots as inorganic phosphate (Pi). Phytates, the major portion of organic phosphorus, often form salts with different ions and are found in less soluble or precipitated forms and hence cannot be utilized by plants. P is a major limiting factor in most agricultural systems. Modern high intensity agriculture is heavily dependent upon external inputs - phosphatic fertilizers being one of them (Chowdhury and Zhang 2021). Modern cereal cultivars, like hybrid maize, require a high dose of externally applied P-based fertilizers. However, it is estimated that more than 90% of the applied P remains unavailable to the crop, and results in environmental pollution via acidification, eutrophication, etc. Intake of phosphate contaminated water causes serious health problems. P is a non-renewable natural resource, mined as rock phosphate and its global reserves may be depleted in 50–100 years (Cordell et al. 2009). Thus, dependence of contemporary agriculture on P fertilizers, poses major food security and sustainability challenge (Udert 2018). Strategies to optimize P use in agriculture needs to be accorded high priority. Improvement of P use efficiency (PUE) in cereal crops could be a major tool to achieve this goal.

Plants have developed a wide range of strategies to adapt to Pi deficiency. These include changes in plant morphology (reduced primary root growth, enhanced number of lateral roots and root hairs); biochemical changes (anthocyanin and starch accumulation), and secretary proteins induction (induction of acid phosphatases, organic acid and RNase secretion). At molecular level, increased expression of Pi transporter genes, differential expression of transcription factors and expression of Pi responsive microRNAs (miRNAs) are also associated with adaptation response to Pi stress (Misson et al. 2005; Nguyen et al. 2015). In the last decade, significant progress has been made in identifying signaling components and their role in responses to Pi starvation in plants (Yuan and Liu 2008). Under Pi deprived condition, the transcriptional regulation of a set of Pi responsive genes occurs through various transcription factors by binding to their respective *cis*-targets present in the promoter region of these Pi responsive genes. Different transcription factors known to be involved in Pi responses include PHR1, ZAT6, WRKY75, WRKY6, MYB26, and BHLH32 in *Arabidopsis*; OsPHR2 and OsPTF1 in rice; ZmPTF1 in maize (Chen et al. 2009; Li et al. 2011) etc. Among these, PHR1 (PHOSPHATE STARVATION RESPONSE1), a MYB domain-containing transcription factor, is one of the most characterized. In recent years, some SPX domain-containing proteins (nuclear proteins) have also been shown to act as important feedback regulators of PHR2. The SPX domain-containing proteins are themselves activated by OsPHR2 under Pi starvation in both *Arabidopsis* and rice (Wang et al. 2009; Liu et al. 2010; Wang et al. 2014). PHR1 is itself regulated post-translationally through the action of SIZ1. SIZ1 is a SUMO E3 ligase that is localized in the nucleus. In *Arabidopsis*, it has been shown that PHR1 is the direct target of SIZ1 and the Phosphate Starvation Induced (PSI) genes were found repressed in *siz1* mutant even in Pi deficiency (Miura et al. 2005). Beside these adaptive responses, acid phosphatase (APase) is also an important participant in mobilization and utilization of organic P under Pi deprived condition (Tran et al. 2010; Tian and Liao 2015). These secreted APases are involved in release of Pi from organophosphates in the plant external environment and increase the availability of Pi to be absorbed by plant root. Several PSI secreted APases have been characterized in vascular plants, like lupin (Ozawa et al. 1995; Li and Tadano 1996; Miller et al. 2001), tobacco (Lung et al. 2008), common bean (Liang et al. 2010), tomato (Bozzo et al. 2006), and *Arabidopsis* (Veljanovski et al. 2006; Wang et al. 2011). In *Arabidopsis*, 11 out of 29 members of Purple Acid Phosphates (*AtPAP*) are transcriptionally up-regulated by Pi starvation (Zhu et al. 2005; Wang et al. 2011).

Among the cereals, maize is the most widely produced and consumed cereal. Owing to its emergence as an industrial and feed crop; with rising worldwide demand, improvement of PUE in maize is likely to have a major impact on sustainability. Identification of key regulatory genes playing pivotal role in acquisition, transportation and utilization of Pi in maize could pave way for greater understanding of this phenomenon in this important crop. In the present study, apart from characterizing the physiological effects of Pi starvation, we have identified homologs of *Arabidopsis* Pi-responsive genes in maize using *in-silico* approaches such as sequence homology and protein functional domain analysis. In addition, expressions of identified Pi responsive genes have been analyzed in root and leaf under Pi starvation. Our study revealed the differential expression of identified Pi-responsive genes belonging to different metabolic pathways under phosphate deprived conditions in maize. To attribute a basis to observed differential expression of the Pi-responsive genes under phosphate deprived conditions, *cis*-regulatory elements present in the upstream promoter region of the selected two genes have also been predicted *in-silico*. Finally, a graphical abstract of transcriptional regulatory model for the Pi starvation responses in maize has been proposed on the basis of present study of selected maize regulatory genes and the previously reported literature (Fig. 1).

**Fig. 1.**
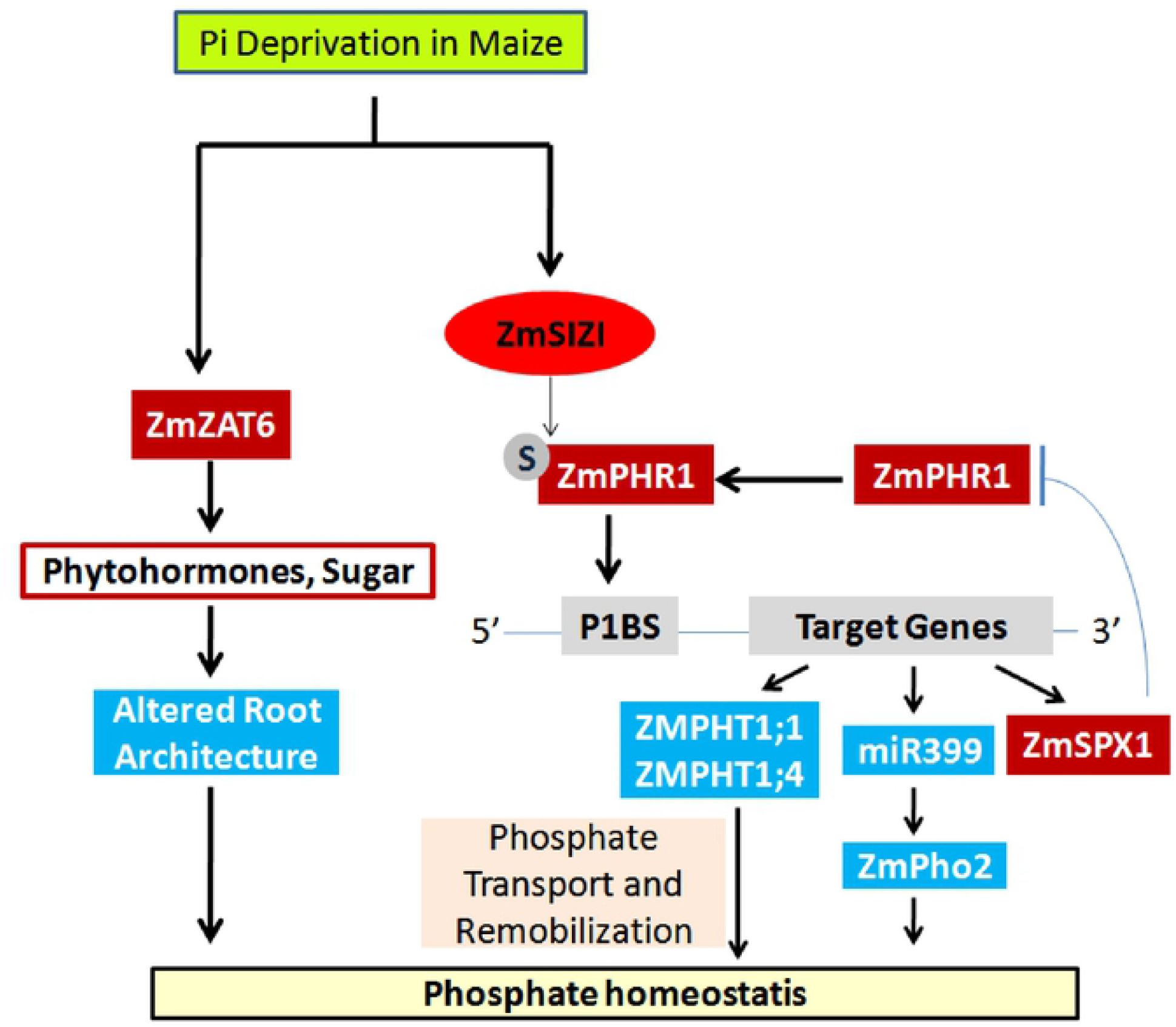
Graphical abstract of the study. Diagram shows regulatory genes and their downstream outcome involved in phosphate deprived condition in maize plant. The hypothesis is derived from sequence and domain similarity of studied maize genes with *Arabidopsis* and their real time expression analysis in shoot and root tissues and the expression similarity with characterized genes. Red colored boxes show regulatory genes and blue boxes denote downstream Pi responsive genes studied in our work. Small gray round circle denotes for sumoylation of PHR1 (phosphate starvation response 1), a transcription factor. P1BS is the cis-regulatory element found on promoter region of many phosphate responsive genes, where PHR1 transcription factor binds and regulate their expression.

## Material and methods

### Plant material and growth conditions

Pi stress tolerant maize (*Zea mays* L.) inbred line HKI-163 was used in this study (Ganie et al. 2015). The seeds were soaked in distilled water for 4 to 6 h and rinsed 4 to 5 times with sterile water followed by surface sterilization by washing with 70% ethanol for 2 min. The seeds were again washed with sterile water (4 to 5 times) followed by treatment with 0.1% HgCl_2_ for 10 min. Seeds were washed again five times with sterile water. The seeds were kept in a sterile germinating paper and then kept for germination at 30±2°C for 4 days in dark. The resulting seedlings were transferred to a high-Pi (1 mM KH_2_PO_4_) nutrient solution and allowed to grow for 2-3 days, after which the seedlings were carefully removed and two sets were prepared for the study. The first set was transferred to modified Hoagland’s hydroponic solution with low Pi (LP) - basal medium + 5 µM KH_2_PO_4_ and 1 mM KCl. The second set was transferred to modified Hoagland’s hydroponic solution with high or sufficient Pi (SP) - basal medium + 1 mM KH_2_PO_4_ (Li et al. 2007; Jiang et al. 2017). Both the sets were allowed to grow for 21 days hydroponically under LP and SP conditions. The nutrient solution was replaced every 3 days. The plants were grown at 25-30°C/ 20 - 25°C (day/night) with a photoperiod cycle of 14 h of light (500-600 µmol m^-2^ s^-1^) and relative humidity of 65% in a phytotron.

### Phenotyping

Six plants from each of the two Pi treatments were harvested after 21 days of treatment and washed with water. The plants were placed in paper bags and dried at 60°C for 48 h. Fresh and dry weights of shoots and roots were determined using an analytical balance (Mettler Scientific, Highstown, NJ). The shoot and root lengths were measured with meter scale. The plants were photographed and the root architecture/ root hair density was studied using phase contrast microscope (Olympus CX41).

### Estimation of plant Pi content

The Pi concentration accumulated in LP and SP grown plants was determined as per Murphy and Riley (1962). Briefly, 10 mg of plant part was crushed with the help of mortar and pestle and 8 ml sterile water was added. A total of 1.6 ml of the mixed reagent (125 ml of 5 N sulphuric acid, 1.5g ammonium molybdate, 75 ml of 0.1 M ascorbic acid solution and 12.5 mg potassium antimonyl tartrate dissolved and made up the volume to 250 ml) was added and diluted to 10 ml volume with water. Standard curve was made by using 0.2µg, 0.4 µg, 0.8 µg, 1.6 µg and 3.2 µg Pi/ml standard Pi solutions (0.1757 g of potassium dihydrogen phosphate in 1 l of sterile water amounting to 40 mg Pi/l) in 10 ml of final volume. The reaction was mixed well. After 10 min, the reaction mixture was centrifuged at 6000 rpm for 5 min. Supernatants were taken in other labeled tubes. Optical density was measured at 882 nm using 1 ml cuvette. Blank reading was determined by freshly distilled water with mixed reagent. Plant Pi content (expressed as µg Pi/mg of dry weight of the plant part) was calculated as per the standard graph.

### Bioinformatic Analysis

Maize genome sequences were downloaded from http://www.maizesequence.org/index.html, and the National Center for Biotechnology Information (NCBI) genome databases. Sequences of *Arabidopsis* genes characterized for Pi responsiveness and the regulatory genes were used as queries to search against the maize protein database with BLASTP program. Hits with Expectation (E)-values below 0.0001 were selected for further domain analysis. All selected sequences from BLAST program were analyzed by functional domain similarity, as confirmed by SMART (http://smart.embl-heidelberg.de) and Pfam (http://pfam.sanger.ac.uk/) databases (Finn et al., 2006; Letunic et al., 2009). The PLACE website (http://www.dna.affrc.go.jp/PLACE/signalscan.html) was used to predict *cis*-elements in the probable promoter regions of Pi responsive genes in the 1000-bp genomic DNA sequences upstream of the initiation codon (Higo *et al*. 1999).

### Primer design and specificity checking

The primers were designed using Primer 3 software provided by NCBI online tools. The specificity of the primer pair sequence was checked against maize genome transcripts (CDS) from the NCBI database using the BLAST program. The primers used are listed in Supplementary Table S1.

### RNA isolation and DNase Treatment

Total RNA was isolated from the root and the leaf samples from LP and SP grown plants harvested at 21 days after treatment (DAT). Approximately, 100 mg of root and leaf samples were crushed in liquid N_2_ using mortar pestle separately. After evaporation of liquid N_2_, 700ul of TRIzol reagent (Thermo Fisher Scientific, USA) was added directly to the mortar pestle and allowed to thaw. After liquefaction of the sample, another 700 ul of TRIzol was added and transferred to a 2 ml centrifuge tube and kept for 2-3 min at room temperature. Further, 300 μl of chloroform was added and the tubes were capped securely and shaken vigorously. The samples were centrifuged at 12000 rpm for 15 min at 4°C. The aqueous phase was carefully transferred to a fresh RNase-free centrifuge tube. RNA was precipitated by adding equal volume of isopropyl alcohol. This mixture was incubated at room temperature for 10 min followed by centrifugation at 12000 rpm at 4°C for 5min. The supernatant was discarded and the pellet obtained was washed with 1 ml of 75% ethanol per 1ml of TRIzol reagent used by tapping only. Subsequently, the samples were vortexed briefly and centrifuged at 12000 rpm for 5 min at 4°C. The RNA pellet obtained was air dried for 5-10 min. The RNA was dissolved in 40 μl RNase-free diethyl pyrocarbonate (DEPC) treated water. About 1 µl of DNase solution (8 µl of 10X DNase buffer, 3U DNase mixed in H_2_O) was added to 10 µl of the RNA sample and kept at 37 °C for 10 min. The digested RNA was column purified before reverse transcription reaction.

### Reverse transcription-polymerase chain reaction (RT-PCR)

The DNase treated RNA was used for the synthesis of first strand cDNA using Superscript III reverse transcriptase kit (Invitrogen, USA). Before performing the RT reaction, 4μg total RNA was mixed with 1 µl of dNTPs and 1 µl of OligodT primer and incubated at 65°C for 5 min. The tubes were immediately transferred on ice and kept for at least 1 min. The cDNA synthesis mix (1X RT buffer, 1.25mM MgCl_2_, 10 Mm dithiothreitol (DTT), 1 U RNaseOUT recombinant ribonuclease inhibitor and 1U SuperScript III RT) was added to the RNA. The reaction was incubated at 50°C for 50 min, followed by heat treatment at 85°C for 5 min. Finally, the samples were treated with 1 µl RNase H at 37°C for 20 min. These cDNA samples were used for quantitative real-time PCR (qRT-PCR) and semi-quantitative RT-PCR analyses.

### Quantitative real-time PCR (qRT-PCR) conditions and analyses

The qRT-PCR was carried out to know the relative transcript levels of the Pi responsive genes in maize in response to high Pi and low Pi conditions treatments. The qRT-PCR was performed using the Brilliant II SYBR Green QPCR Master mix (Agilent) in real time PCR (Agilent Technologies, USA) detection system. A 20 μl reaction-mixture containing 10µL of SYBR^®^ Green premix, 0.25 µM of each primer pair (Supplementary Table S1) and 2.0 µl cDNA template was gently mixed in 96-well Real-Time PCR plate, and centrifuged at 200 rpm for 2 min to spin down all reaction components in the plate. The reactions were carried out with the following thermal profile: 50°C for 2 min, 95°C for 3 min, followed by 40 cycles of 95°C for 10 s, 60° C annealing for 20 s. After 40 cycles, the specificity of the amplifications were tested by heating from 65°C to 95°C with a ramp speed of 1.9°C/min, resulting in melting curves. The reference control genes were measured with three replicates in each PCR run, and their average cycle threshold (CT) was used for relative expression analyses. The *Actin* gene from maize was used as reference gene to normalize the expression values. The log2 fold change value was calculated based on 2^−(ΔΔCt)^method.

### Semi-quantitative RT-PCR

The optimum numbers of cycles for semi-quantitative RT-PCR were worked out by using real time PCR amplification curve, so that optimum difference between control and treated could be visualized in semi-quantitative PCR amplification. For semi-quantitative analysis, PCR amplifications were performed in 25 µl of total volume reaction containing 100 ng template cDNA, 1X Taq Polymerase buffer, 1.5 mM MgCl_2_, 1 mM dNTPs, 0.4 µM of each forward and reverse primer pair and 2U of Taq DNA polymerase. The PCR amplification was achieved in a BioMetra thermal cycler with amplification conditions of 94°C for 3 min (1 cycle) followed by 30 cycles of 94°C for 30 s; 60°C for 30 s; 72°C for 30 s and finally 72°C for 5 min extension. PCR product (10 µl) was analyzed by electrophoresis on 3% agarose gel along with 50 bp DNA ladder. The bands were detected by ethidium bromide staining and photographed under UV light using the Gel Documentation system (Alpha Innotech).

## Results

### Effect of Pi starvation on plant growth and Pi accumulation

After 21 days of phosphate treatment, the plants grown under LP conditions depicted observable symptoms of Pi starvation with respect to shoot growth, root architecture, lateral roots, root hairs, leaf coloration and stems (Fig. 2 and Table 1). Under Pi starvation, the shoot fresh and dry weight decreased by 33.2 and 29.9%, respectively, while the root fresh and dry weight decreased by 26.1 and 8.1%, respectively, corroborating the role of Pi in biomass accumulation. The overall plant fresh and dry weight reduced by 29.9 and 26.8%, respectively. The shoot length and stem girth were reduced by 14.7 and 21.0%, respectively. There was a pronounced increase in root length due to Pi starvation. The average root length increased significantly by 85% (P <0.05). Also, as discerned by phase-contrast microscopy (Fig. 2e), there was a proliferation of root hairs in the roots of Pi starved plants. Pi starvation also led to lesser number of crown roots (Fig. 2c; Table 1) and lengthier lateral roots (Fig. 2d). Further, the Pi stress treatment was confirmed by measuring the levels of endogenous Pi content in leaves and roots of the plant grown under Pi sufficient and deficient conditions. Under Pi stress (LP) conditions, both shoots and roots accumulated < 0.1 µg Pi mg^-1^of dry weight biomass, in contrast to accumulation of approximately 1 µg Pi mg^-1^ of dry weight biomass under Pi sufficient (SP) conditions (Fig. 3). This confirms the efficacy of the hydroponic experiment in imposing quantifiable Pi stress on the plants.

**Table 1.**
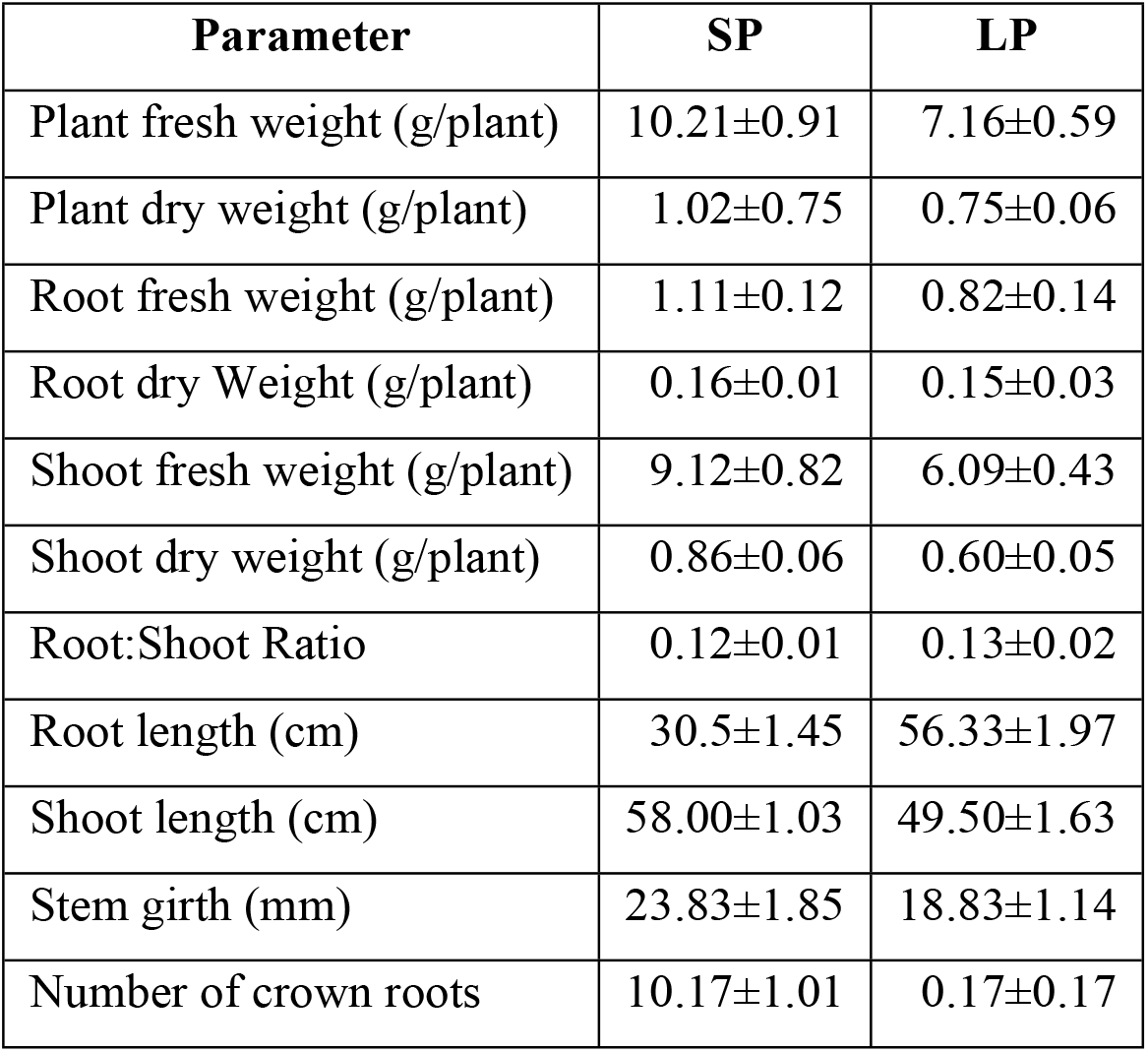
Physiological parameters of maize genotype HKI 163 grown under low phosphate (LP) and sufficient phosphate (SP) conditions for 21 days. All the values shown here represent the mean± SD of six plants.

**Fig. 2.**
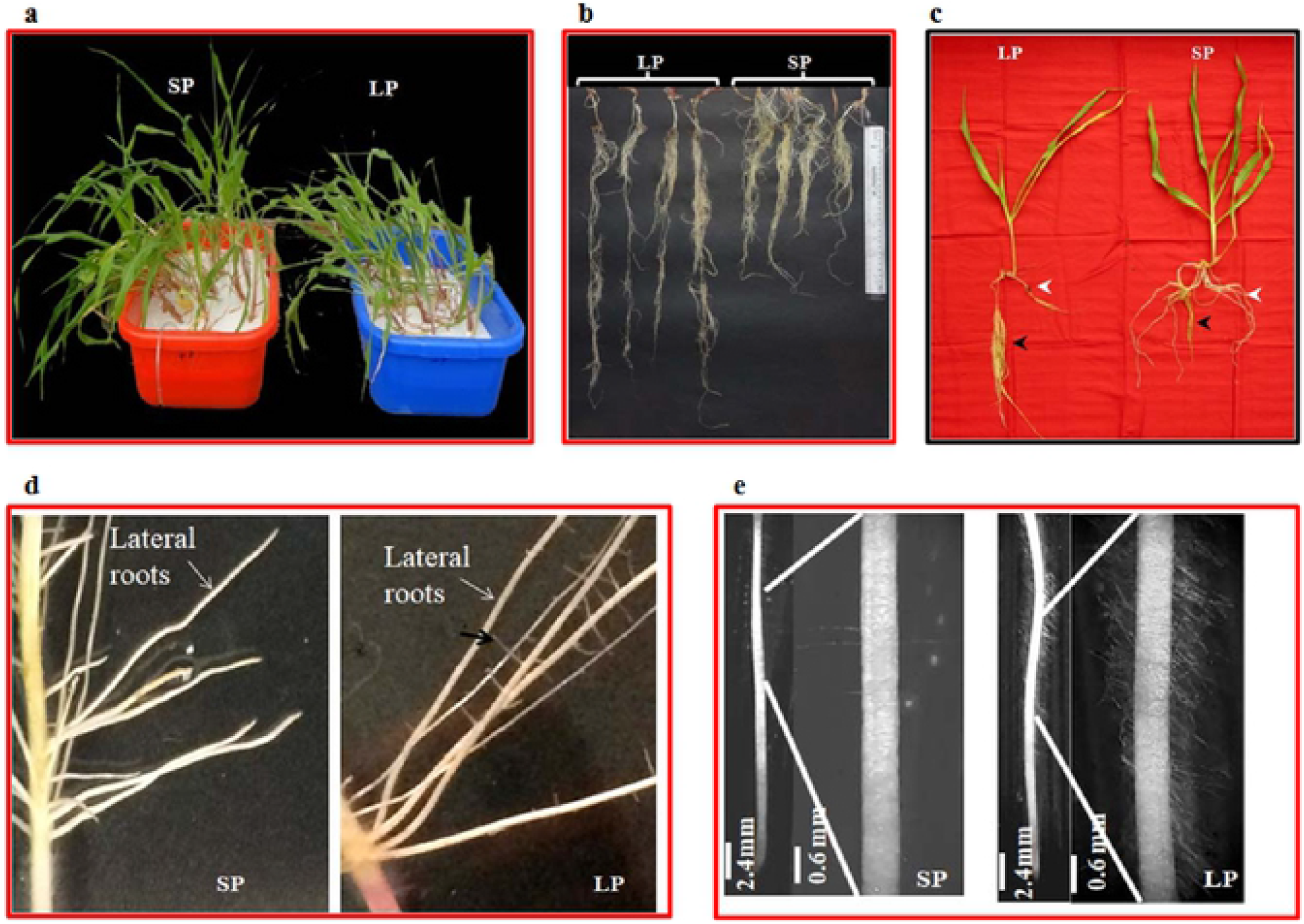
Effect of phosphate availability on plant growth and root architecture in a phosphate stress tolerant maize genotype HKI-163 grown in two growth conditions-deficient (LP; 5 uM phosphate) and sufficient (SP, 1mM phosphate). (**a)** Plant growth under hydroponics in SP and LP conditions. **(b)** Root length under SP and LP conditions. **(c)** Root architecture, Black arrow: main root with lengthy lateral roots (in LP condition) and less lengthy lateral roots (in SP condition). White arrow: crown roots, more in numbers under SP condition and less in LP. **(d)** Close-up of roots under LP and SP treatment. White arrow: main root with lengthy lateral roots (in LP condition) and less lengthy lateral roots (in SP condition). **(e)** Phase contrast microscopy images of root hairs-main root with short and less root hairs (in SP condition) and lengthy and more root hairs (in LP condition). Photographs were taken after 21 days of treatment.

**Fig. 3.**
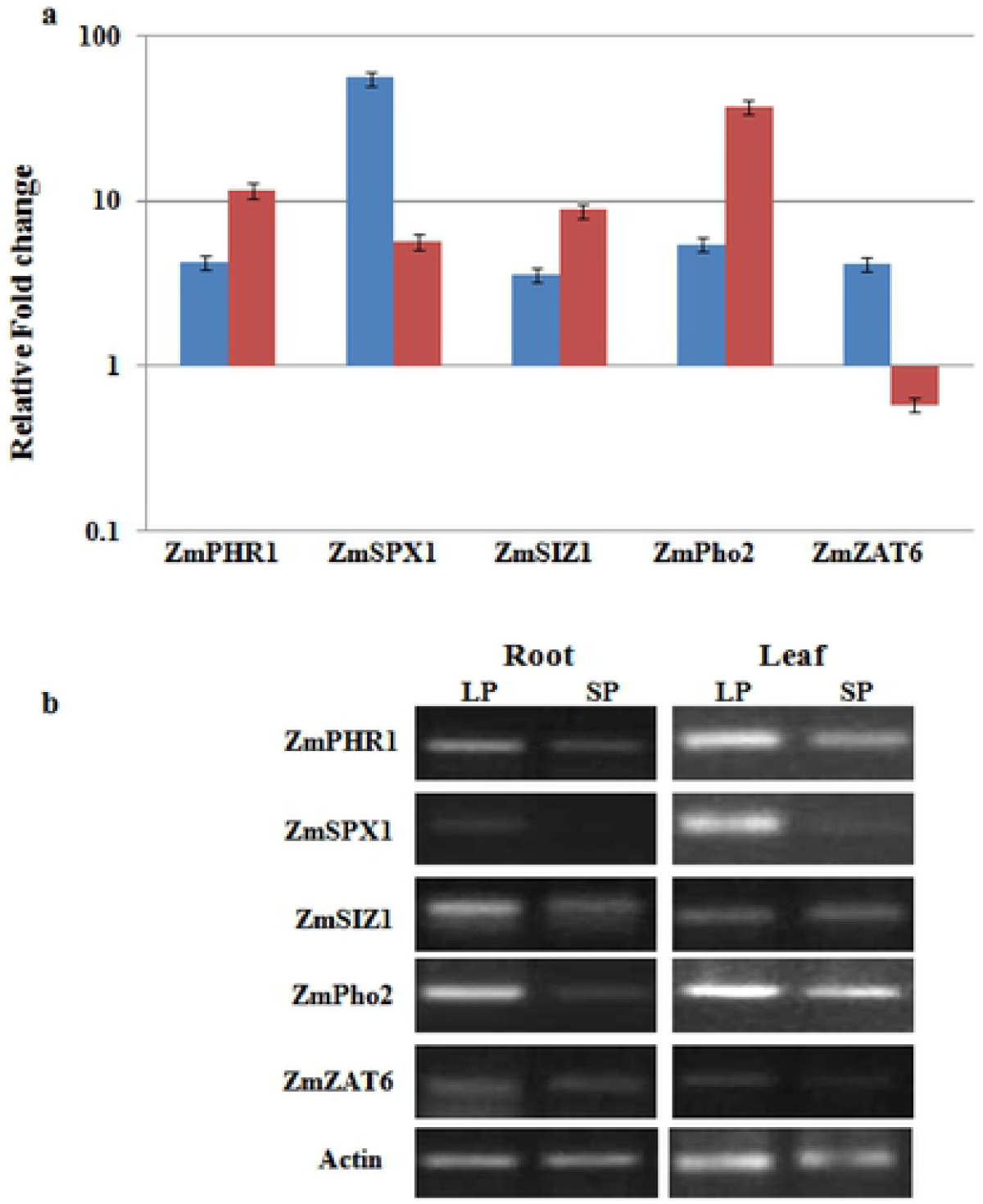
Effect of phosphate availability on maize Pi accumulation. To determine the phosphate content of plants, plants were harvested, dried, and total phosphorus was assayed by method described by Murphy and Riley. Total Pi content is given in µg Pi/mg of dry weight of the plant part. Values shown represent the mean of six plants and triplicate sample from each sample. LPR, Maize root under Low Phosphate condition; SPR, Maize root under Sufficient Phosphate condition; LPL, Maize leaf under Low Phosphate condition; SPL, Maize leaf under Sufficient Phosphate condition.

### Comparative sequence and domain analysis of potential Pi-starvation responsive genes in maize

For identification of potential Pi-responsive genes, three lines of evidences were relied in this study. First, a comparative sequence analysis was carried out to identify potential maize homologs of the reported Pi-responsive genes from *Arabidopsis thaliana* with 52 to 93% sequence similarity and up to 99% query coverage. Utilizing minimum E-value criteria, most of the selected genes from *A. thaliana* had clear homologs in *Z. mays* (Supplementary Table S2). Secondly, the isolated sequences were subjected to Pfam and SMART databases for functional domain analysis. We found that the corresponding maize genes had largely the same domain architecture, position wise and structure wise, as seen in their counterpart in *Arabidopsis* (Supplementary Fig. S1-S3). The genes could be grouped into three major classes on the basis of their functions, *viz*., the genes involved in regulation; the major secretory proteins; and the Pi transporters. Four major regulatory genes were included in the analyses. The *PHR1* gene is a transcription factor involved in regulating a subset of the genes which are responsive to Pi starvation (Rubio et al. 2001; Bari et al. 2006). The maize *PHR1* gene has the myb-DNA binding domain like *Arabidopsis*, in between 200 to 400 amino acid residues at C terminal of the protein. The maize SPX1 domain containing protein, also has the same domain architecture as AtSPX1 protein at amino terminus, with only difference that the domain size was slightly larger as compared to AtSPX1. SIZ1, a E3 SUMO-protein ligase has domain architecture like their counterpart in *Arabidopsis* i.e. presence of all the functional domains *viz*. the SAP domain, the DNA-binding domain, the plant homeodomain (PHD), a ubiquitin ligase and Zinc finger (Znf) domain containing multiple finger-like protrusions for salt bridge formation. Another important regulatory protein ZmPho2-an ubiquitin-conjugating enzyme E2, catalytic (UBCc) domain containing protein was found identical to corresponding *A. thaliana* protein at C terminal. ZAT6-the Zinc finger domain containing transcription factor was also found to possess same domain architecture (Supplementary Fig. S1). Among the secretory proteins, PAP10 and PAP17 possessed the entire metallophos domain at proper position as AtPAP10 and AtPAP17 with only a slight difference in PAP17 metellophos domain. The maize PAP2 and ZmRNS1 had the same SANT domain and ribonuclease_T2 domain respectively as in AtPAP2 and AtRNS1 (Supplementary Fig. S2). Both the high affinity transporters (the MFS: major facilitator superfamily) studied had the similar sugar transporter-like domain and 12 trans-membrane domains as other MFS transporters (Supplementary Fig. S3). The third and the final line of evidence towards identification of Pi starvation induced genes were based on actual transcript expression analysis of the genes, as presented further.

### Expression pattern of regulatory genes involved in Pi starvation response in a Pi stress tolerant maize genotype

In order to confirm the genes identified by sequence homology and domain analysis, semi-qRT and real-time PCR analyses were conducted in parallel to verify the validity of these genes. Five regulatory genes that play central role in the Pi homeostasis-*ZmPHR1, ZmSPX1, ZmSIZ1, ZmPho2* and *ZmZAT6* were selected for further validation. These genes were chosen because they had been previously implicated to be involved in regulatory pathways modulating Pi starvation response, or because of their possible contribution to Pi homeostasis in plants under Pi deficiency (Rouached et al. 2010). The transcription levels of mRNA were significantly higher for all the genes at LP conditions compared with SP (control) conditions in both root and leaf, except *ZAT6*, which was specifically up regulated in leaves only. Real-time PCR analysis (Fig. 4) revealed that under Pi starvation, *ZmPHR1, ZmSPX1, ZmSIZ1, ZmPho2* and *ZmZAT6* genes were 4.28, 55.01, 3.57, 5.41 and 4.18 times overexpressed in the leaf. In the roots, the *ZmPHR1, ZmSPX1, ZmSIZ1, ZmPho2*genes were 11.49, 5.64, 8.76, 36.79 times over expressed, while *ZmZAT6* was under expressed 0.58 times. The above results also correlate to the band intensities of the respective amplicons as visualized in the semi-qRT-PCR.

**Fig. 4.**
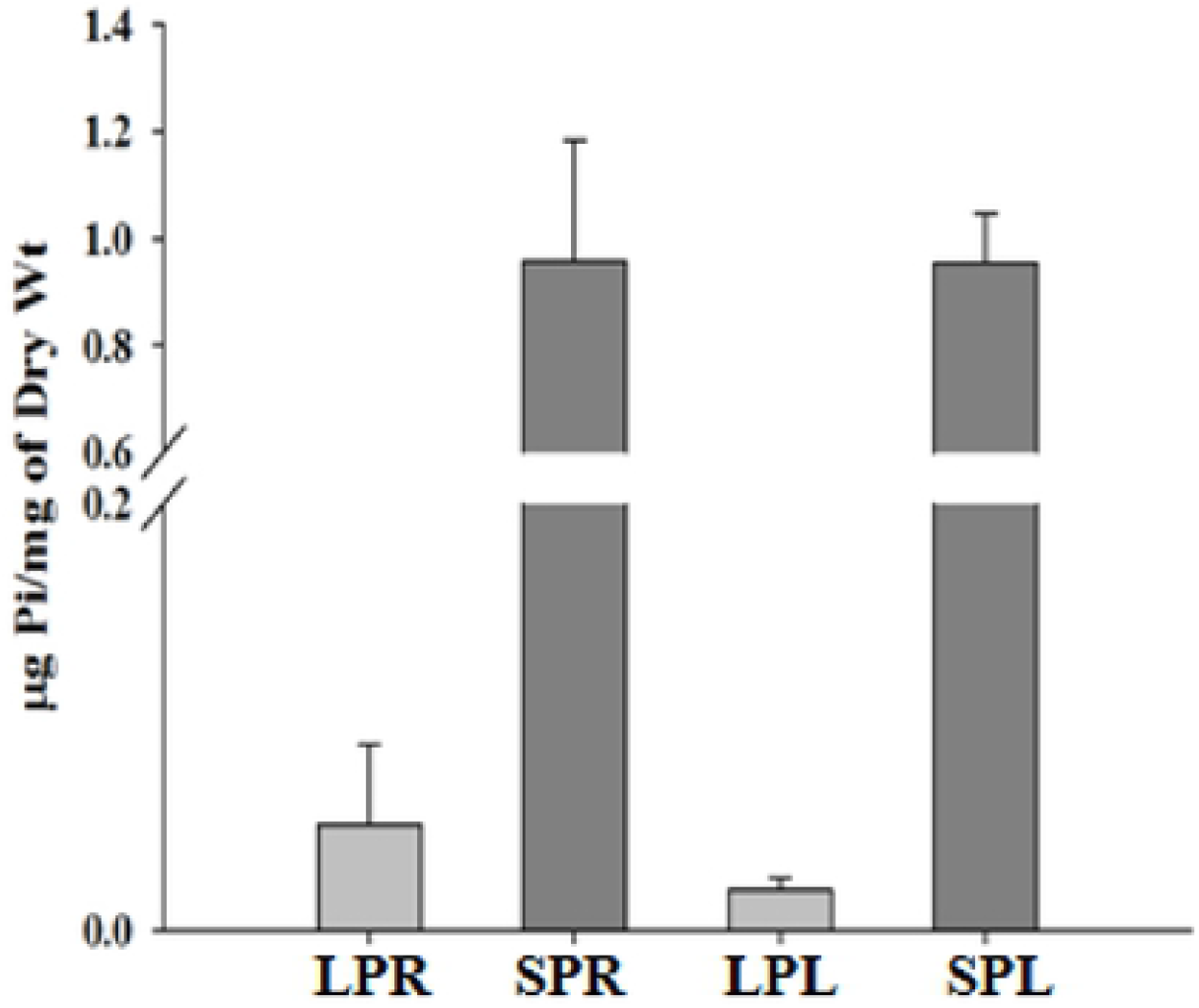
Expression analysis of regulatory genes involved in phosphate responsive pathway. **(a)**The qRT-PCR based expression analyses of predicted regulatory genes at phosphate deprived conditions in HKI-163 inbred line of maize grown under phosphate deprived (5 µM KH_2_PO_4_) and sufficient phosphate conditions (1 mM KH_2_PO_4_). *Y*-axis represents the relative fold change values at low phosphate condition as compared to respective genes at sufficient phosphate conditions. **(b)** Semi-quantitative RT-PCR expression analyses using same set of primers at 30 PCR cycles of same genes for re-confirmation.

### Pi starvation mediated modulation of PAPs and RNase genes involved in P solublization and remobilization

As an initial characterization of gene expression in maize plants, an expression pattern of 4 selected genes (*ZmPAP2, ZmPAP10, ZmACP5* and *ZmRNS1*) were examined using qRT-PCR and RT-PCR in leaf and root tissues (Fig. 5). The differential/varied expression levels of these genes were found in the leaves and roots as expected, except *PAP10* gene which was down regulated in the roots. As expected, the expression of *ZmPAP2* was slightly upregulated in both leaf and root tissues. The expression level of putative *PAP10* was slightly high (2.97 fold) in leaf but extremely low (0.11 fold) in root tissue at Pi deprived condition as compared to sufficient Pi condition. RT-PCR showed that both *ZmACP5* and *ZmRNS1* mRNA levels were strongly upregulated in the root tissue (74.40 and 115.09 fold respectively). As expected, the expression level of *ZmACP5* in leaf tissue was high, while the *ZmRNS1* was down regulated in leaves.

**Fig. 5.**
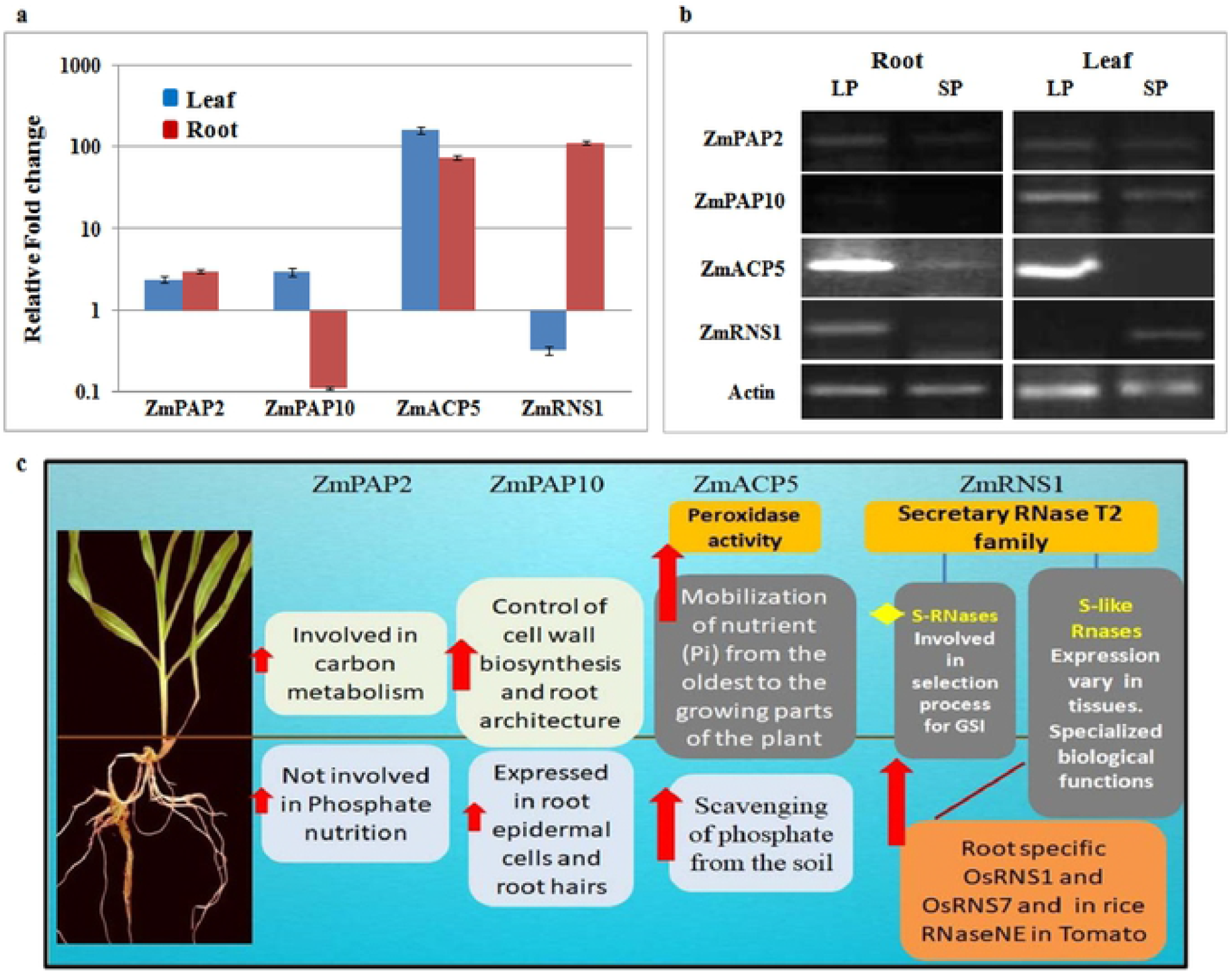
Expression analysis of purple acid phosphatases (PAPs) and ribonuclease (RNase) genes involved in phosphate responsive pathway. **(a)** The qRT-PCR based expression analyses of predicted PAPs and RNase genes at phosphate deprived conditions. *Y*-axis represents the relative fold change values at low phosphate condition as compared to respective genes at sufficient phosphate conditions. **(b)** Semi-quantitative RT-PCR expression analyses using same set of primers at 30 PCR cycles of the same genes for re-confirmation. **(c)** A schematic representation of function and expression on known PAPs and RNase. Red arrow mark shows expression of respective known genes.

### Modulation of high affinity Pi transporters

Particularly high affinity Pi transporters are expected to perform crucial functions in Pi acquisition and remobilization at Pi deficiency. With a particular interest in these genes, we chose two genes (*ZmPht 1;1* and *ZmPht 1;4*) on the basis of sequence similarity with *A. thaliana* and the domain analysis. Both these genes were upregulated in leaves and roots under Pi deprived condition with varied level of expression. *ZmPht1;4* showed a higher level of expression in root tissues (fold expression in root was 22.61 fold whereas 5.34 fold in leaf) whereas *ZmPht1;1* was expressed more in leaf tissues (169.55 fold expression in leaf and 30.45 fold in roots) in response to Pi stress (Fig. 6).

**Fig. 6.**
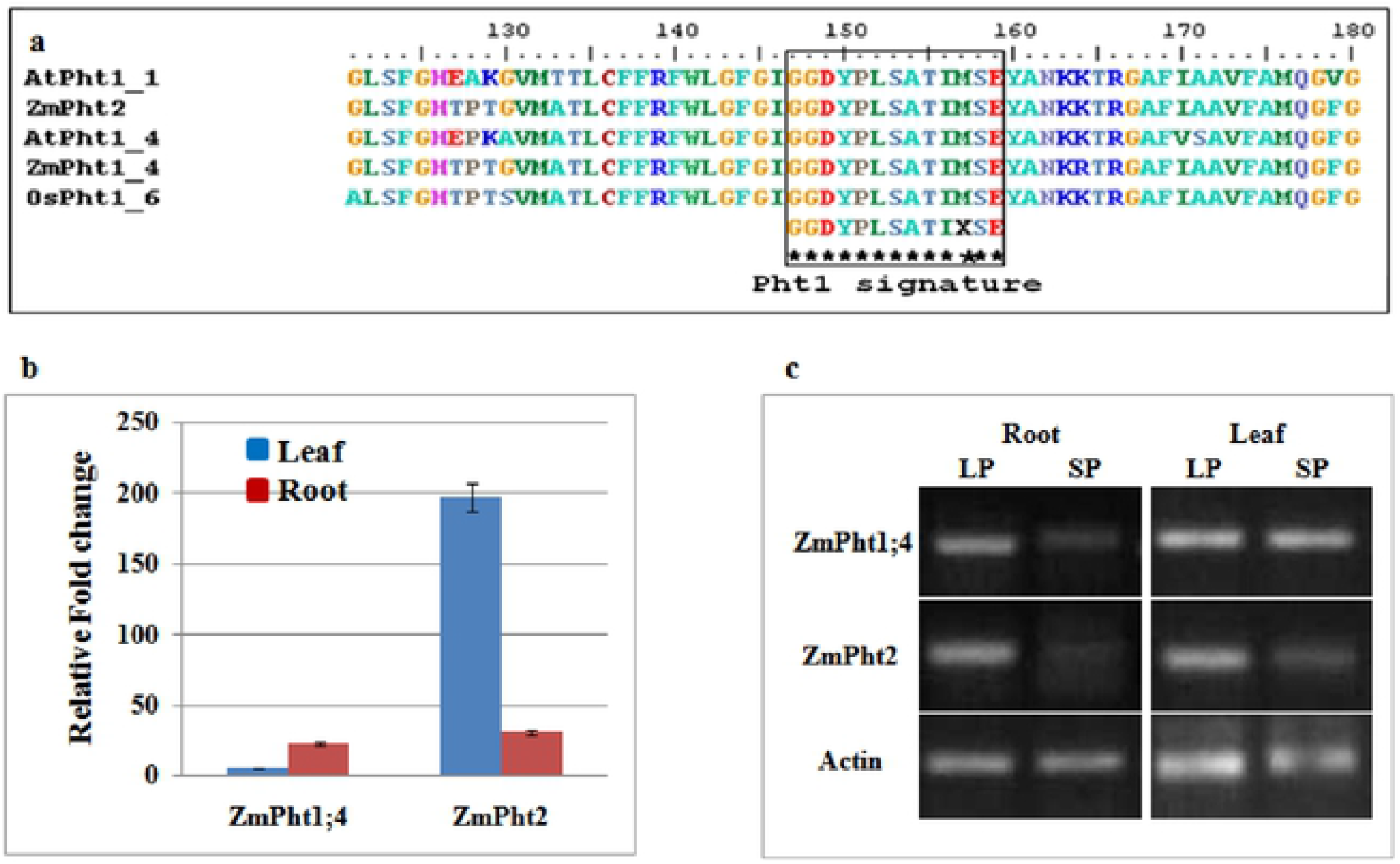
Expression analysis of high affinity Pi transporters in maize. **(a)** Alignment of the peptide sequences of known high affinity Pi transporters from *Arabidopsis*, and rice with identified high affinity phosphate transporters of maize; identical peptides are highlighted in the same color. The *Pht1* signature sequence is shown in the box. **(b)** Semi-quantitative RT-PCR expression analyses using the same set of primers at 30 PCR cycles of the same genes for re-confirmation. **(c)** The qRT-PCR based expression analyses of the predicted high affinity Pi transporters genes at phosphate deprived conditions. The *Y*-axis represents the relative fold change values at low phosphate condition as compared to respective genes at sufficient phosphate conditions.

## Discussion

Maize is a widely cultivated grain crop and a major source of feed, food and industrial raw material. The identification of Pi starvation-induced genes and delineating the fundamental molecular-genetic mechanisms in Pi stress response are important steps towards engineering high PUE in this crop. A few regulatory genes and a countless other Pi responsive genes have been identified as adaptation components of Pi starvation in plants, especially in model plants like *Arabidopsis* (Lopez-Arredondo et al. 2014). However, identities and functional biology of majority of Pi stress responsive genes in maize, especially so in the tropical and the sub-tropical genotypes, remained sketchy. In this study, we chose a tropically adapted maize inbred line (HKI-163) that is tolerant to Pi stress and determined its physiological and molecular response to Pi starvation under controlled conditions.

### Plant growth characteristics under Pi stress

Roots are the tissues involved in Pi uptake in crop plants. Root growth and morphology is dramatically changed when Pi is deprived in root zone. These changes lead to increase in root length, increase in the root to shoot ratio and lateral root length. All these measures are the adaptive response to Pi deficiency. The relationship between Pi assimilation efficiency and root morphology has been analyzed in maize (Peret et al. 2011). Consistent with these findings, our studies demonstrate increased root length, reduced total biomass, proliferation in root hair, increase in the length of the lateral and nodal roots, increased root: shoot ratio and reduced level of total Pi uptake in root and shoot significantly (P < 0.05) at low Pi condition compared to sufficient Pi condition (Fig. 2, 3 and Table 1). Our physiological data with respect to Pi starvation were consistent with previous studies on maize plants subjected to low Pi stress for 17 days (Li et al. 2007). The HKI-163 inbred line has been reported to exhibit higher shoot dry weight, total plant biomass, root length, root dry weight, leaf area, total P uptake and PUE as compared to Pi stress sensitive maize genotypes (Ganie et al. 2015).

### Genes involved in regulation during Pi stress

At a molecular level, Pi deficiency is regulated both at the transcriptional and post-transcriptional levels. So far, the major actor coordinating these various regulations is PHR1, *via* the PHR1-PHO2–miRNA399 pathway, conserved amongst flowering plants. Beside this transcription factor, few other regulators have also been identified including ZAT6, WRK75, bHLH32 and MYB62 (Yi et al. 2005; Chen et al., 2007; Devaiah et al. 2009). In this study, we selected homolog of PHR1, SPX1, SIZ1 and PHO2-genes contributing in PHR1 regulation pathway and another less studied TF, ZAT6. Study of these genes with their sequence similarity with *Arabidopsis*, functional domain analysis and finally expression analysis in root and leaf tissues by semi-quantitative RT-PCR and quantitative real-time PCR (qRT-PCR) reveal that these selected accessions might be involved in Pi stress response in maize and might work as previously described regulators.

From the maize genome, we took a highly similar sequence homolog to *Arabidopsis PHR1* gene followed by domain analysis (Supplementary Table S2 and Supplementary Fig. S1-S3). Interestingly, the rice genome contains two *PHR1*-like genes, both reported as involved in Pi starvation (Zhou et al. 2008). In the present study, we also found two *PHR1* like gene sequences in maize but we selected only one highly similar sequence for further study. The identified ZmPHR1 had 62 % sequence similarity with AtPHR1. Domain analysis showed MYB domain at C terminal end of the protein as previously characterized in *Arabidopsis* PHR1 and the real time expression was observed as expected. MYB-like domain of PHR1 binds to a DNA motif GNATATNC, termed P1BS (Rubio et al. 2001), which is present in the promoter of many Pi starvation-induced genes and regulate their expression (Franco-Zorrilla et al. 2004; Mission et al. 2005; Müller et al. 2007). Furthermore, it has been shown that PHR1 affects expression of miRNA399 and consequently expression of PHO2 which is involved in Pi homeostasis (Fujii et al. 2005; Aung et al. 2006; Bari et al. 2006; Chiou et al. 2006). So *Pho2* is an indirect target of PHR1 TF via miRNA399. Bioinformatically, we analyzed the miRNA 399 target site on *ZmPho2* gene and P1BS motif on the probable promoter region of miRNA 399 conjecturing that *ZmPho2* might be the indirect target for PHR1 and may have possible role in PHR1 signaling. However, our real time expression data did not support this. So it might be possible that there is another *Pho2* like element present in maize or some other sequence in the maize genome that works as *Pho2*.

Besides *Pho2*, SPX1 is direct target of PHR1. In the presence of Pi, SPX1 displays high binding affinity to PHR1 and sequesters it, so that binding of PHR1 to its target genes via P1BS is inhibited, and their transcription (including that of SPX1) is just at basal level. While in the absence of Pi, the affinity of SPX1 to interact with PHR1 is reduced, so PHR1 is free to interact with its targets, resulting in induced expression of the target genes including SPX1 (Puga et al. 2014; Yao et al. 2014; Zhang et al. 2016). As a result, there is increased expression of SPX1 at low Pi condition. High SPX1 protein levels allow rapid shutdown of PHR1 target gene expression after Pi re-feeding but at low Pi, the affinity of SPX1 is very low, while at constant Pi stress there is high level expression of SPX1 but in inactive form (Puga et al. 2014). Evidence for the importance of the P1BS motif on the promoter region of PHR1 targeted genes has been observed in monocot species (Schünmann et al. 2004). In the present study, beside the sequence similarity and domain analysis of the ZmSPX1 protein, the P1BS motif GNATATNC (PHR1 binding site on the target gene promoter) was also found on the −151 bp position of the ZmSPX1 promoter (Supplementary Table S2, Supplementary Fig. S1 and Fig. 7). So it is a good indicator that the selected sequence encode ZmSPX1 gene. The expression pattern of the ZmSPX1 gene correlates with ZmPHR1 expression data in both root and leaf (Fig. 4). Hence, our study is in agreement with previous studies in *Arabidopsis* with respect to its expression pattern (Puga et al. 2014).

**Fig. 7.**
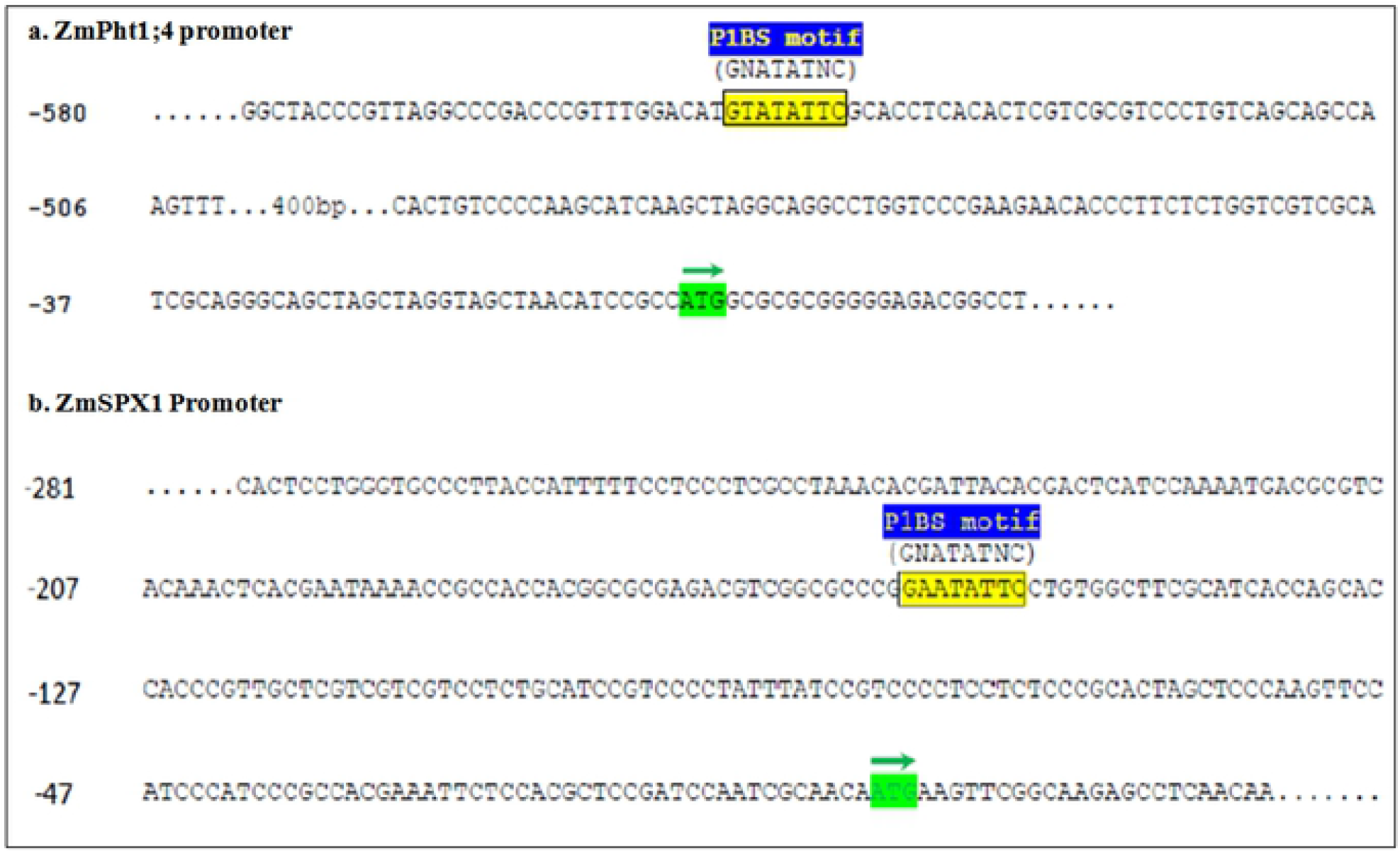
ZmPht1;4 and ZmSPX1 promoter containing P1BS motif required for ZmPHR TF binding. 1000 bp region upstream to the ATG start site of ZmSPX1 gene was scanned for the presence of P1BS (GNATATNC) motif, a SPR1 binding site, by using a MEME online motif finder software. The P1BS motif were found at −542 and −151 bp of the promoter of ZmPht1;4 and ZmSPX1 respectively. Translational start site (ATG) is shown as green highlighted, ATG and green arrow indicates direction of translation. P1BS motif is shown as yellow colored box.

Under low Pi conditions, PHR1 TF is sumoylated by SIZ1, a SUMO E3 ligase that is localized in the nucleus, and this post-translational modification is likely important for PHR1 activity. Sumoylation of PHR1 controls expression of Pi starvation-responsive genes (Miura et al. 2005), although the mechanism of this regulation is still unknown. So the expression of PHR1 and its target genes should be directly correlated with the expression of SIZ1 gene. The up-regulation of the expression pattern of the ZmSIZ1 gene in both root and leaf tissue in the present study also supports this in maize (Fig. 4). Pi stress is known to trigger a significant increase in SUMO-protein conjugate levels (Miura et al. 2005, Kurepa et al. 2003; Miura et al. 2007). Recently, Miura et al. (2005) reported that SIZ1 is also involved in the regulation of root growth in response to Pi starvation.

Beside, PHR1 microarray analysis revealed that ZAT6, a Cys-2/His-2 (C2H2) zinc finger transcription factor, has also been implicated in the regulation of Pi starvation responses and strongly induced during Pi deprivation (Rubio et al. 2001). This gene plays a vital role in seedlings by regulating growth and Pi homeostasis. ZAT6 is a gene induced during Pi starvation that responds rapidly and specifically to the altered Pi status of plants (Devaiah et al. 2007). In the present study, we observed increased expression of ZmZAT6 in maize leaf (Fig. 4) that is in agreement with the *Arabidopsis* microarray data. On the other hand, its reduced expression in roots as the plants grow older allows the primary roots to elongate (Devaiah et al. 2007). In the present study, the expression of the same gene was slightly reduced in root tissue at 25 days of Pi stress (Fig. 4). So the present data in root tissue at 25 days suggest that this gene is reduced to allow root elongation at low Pi at this stage and participate in the root development whereas an increased expression in the leaf suggest its role in Pi-homeostasis in the shoot part. These selected Pi responsive genes make two different regulatory pathways, first PHR1 regulated and second ZAT6 regulated mechanisms for Pi homeostasis and Pi management under Pi stress condition.

### High affinity Pi transporters in maize

Pi uptake by plants operates via two systems, high and low affinity Pi transporter system (Furihata et al. 1992). The high affinity transport system is largely mediated under Pi deprived conditions by plasma membrane-localized Pi transporters belonging to the *Pht1* family (Raghothama 2000). PHT1 belongs to major facilitator super family (MFS) proteins, encoding high-affinity H^+^/Pi co-transporters. It has been previously reported that PHT1 Pi transporters play a critical role in Pi acquisition from soil solution and Pi remobilization within the plant (Nussaume et al. 2011; Gu et al. 2016). In the present study, *ZmPht1;1* and *ZmPht1;4* sequences were found to share a high level of similarity to known *Arabidopsis* and rice high affinity Pi transporters and also have the GGDYPLSATIxSE signature sequence (Fig. 6a). Promoter scan by using MEME (Lescot et al. 2002) revealed that Pht1;4 transporter had the P1BS motif GNATATNC (PHR1 binding site on the target gene promoter) at −542 (Fig. 7a) whereas the promoter of *ZmPht2* does not have any P1BS motif which indicated that the expression of these transporters might be PHR1 dependent and independent respectively. *ZmPht1;4* might be regulated by PHR1 transcription factor in a similar manner as described previously for known high affinity Pi transporters (Bustos et al. 2010; Nilsson et al. 2010; Oropeza-Aburto et al. 2012). Functional characterization shows that some of the *Pht1* members, such as *AtPht1;1, AtPht1;4* and *OsPht1;6* are high-affinity transporters while others are low-affinity transporters, such as *OsPht1;2* (Shin et al. 2004; Ai et al. 2009). Previously, it was confirmed that *HvPht1;1, CmPht1* and a number of *Pht1* family transporters are strongly expressed in the root, and are induced by Pi starvation (Liu et al. 1998; Ai et al. 2009; Jia et al. 2011; Wu et al. 2011; Liu et al. 2014). Like *AtPht1;1, HvPht1;1* and other root specific high affinity transporters, *ZmPht1;4* also exclusively expressed in the root tissues (Fig. 6b, c) suggesting that they might play important role in Pi acquisition from soil. Whereas, *HvPht1;6* and other transporters show enhanced expression in both leaves and roots (Huang et al. 2008); localize in the leaf phloem tissue; and are involved in Pi re-translocation (Rae et al. 2003). Like *HvPht1;6, ZmPht1;1* (*ZmPht1;4*) were expressed in both root and leaf (Fig. 6b, c) suggesting that ZmPHT1;1 is a high affinity transporter that might be participating in Pi re-translocation when Pi is limited in the leaf tissues.

### APases and RNAses involved in solublization and remobilization of Pi during Pi stress in maize

Plant Apases (EC 3.1.3.2) are hydrolase enzymes which catalyze Pi from a group of phosphomonoesters and anhydrides (Duff et al. 1994). Among the Apases, purple acid phosphatase (PAP) has distinctive character of producing purple or pink color in aqueous solution and the presence of seven invariant residues in five conserved metal ligating motifs (Bozzo et al. 2006; Matange et al. 2015; Tian and Liao 2015). Despite low homology between PAPs from different kingdoms, five conserved motifs have been identified, including DXG, GDXXY, GNH (D/E), VXXH, and GHXH (Flanagan et al. 2006; Schenk et al. 2013). Studies show that plant PAP members are involved in P scavenging, recycling and utilization of different forms of extracellular organic P under conditions of P deficiency (Wang et al. 2011; Robinson et al. 2012; González-Muñoz et al. 2015; Liu et al. 2016). Sequence alignment using COBALT, a Constraint-based Multiple Alignment Tool, showed all the 5 PAPs conserved domain in selected maize PAPs (Supplementary Fig. S4). In this study, we have selected two PAPs one is *AtPAP10* and second *AtPAP17* (*AtACP5*) on the basis of sequence, domain and motif similarity. It has been shown that *AtPAP10* mRNA levels were increased 5- and 4-fold in Pi-starved leaves and roots of *Arabidopsis*, respectively (Wang et al. 2011). This suggests that ZmPAP10 may function like AtPAP10 when plants are stressed by Pi deficiency. Pi starvation induces *AtACP5* not only in roots but also in aerial parts of the plant. AtACP5 displayed two type of activities; peroxidase activity and phosphatase activity. Pozo et al. (1999) suggested its probable role in recycling of the Pi from the Pi ester pool of the plant and its participation in the scavenging of Pi from the soil. Similarly, in the present study, the strong over expression of ZmACP5 in both root and leaf under low Pi conditions (Fig. 5) suggests its role in Pi recycling in shoot and Pi scavenging in root surface. AtPAP2, a MYB family protein participates in flavonoid biosynthesis in *Arabidopsis* plant organs (Borevitz et al. 2000). *Arabidopsis* plants overexpressing PAP1 or PAP2 show intense purple pigmentation in many vegetative organs throughout development, and more detailed analysis of PAP1 over-expressing plants shows that some flavonoid biosynthetic genes are expressed constitutively, and the accumulation of anthocyanins is markedly enhanced (Borevitz et al. 2000; Tohge et al. 2005). Previously, it was shown that AtPAP2 is involved in carbon metabolism but not in phosphorus nutrition and it expresses under Pi deprived conditions in both root and shoots (Feng and Lim 2011; Sun et al. 2012). Our results also show slight up regulation of *ZmPAP2* in both root and leaf.

Ribonucleases (RNases) found in cellular compartments of secretory pathway belong to the RNase T2 family of endoribonucleases. These RNases are secreted directly from the cell to produce phosphomonoesters by RNA degradation (Desspande and Shankar, 2002). T2 families of RNases are of two types in plants, S-RNases and S-like RNases (Fig. 5c). The S-RNases are involved in selection process for gametophytic self-incompatibility and expressed in the gametic tissues only (Roalson and McCubbin, 2003). S-like RNases are expressed in many different organs or tissues to achieve specialized biological functions under different stresses including Pi starvation, pathogen infection, wounding and different biotic and abiotic stress. In the present study, a particular RNS was strongly expressed in the root tissue (115 fold expression in low Pi condition as compared to sufficient Pi) but not in the leaf tissues suggesting that it is a root specific RNS (Fig. 5a, b). Transcript level of RNS1 and RNS2 in *Arabidopsis*, RNaseNE in *Nicotiana alata* and RNaseLX and RNaseLE in tomato are induced during Pi limitation (Taylor et al. 1993; Bariola et al. 1994; Kock et al. 1995; Kock et al. 2006). RNaseNE mRNA is expressed in the roots, but not in vegetative tissue (Dodds et al. 1996). Tissue specific expression analysis of RNase T2 transcripts by RT-PCR revealed the root specific expression of *OsRNS1* and *OsRNS7* and show that these two RNases are specifically expressed in roots (MacIntosh et al. 2010). So, ZmRNS1 is an S-like RNase in maize exclusively expressed in root tissue under Pi deprived conditions.

## Conclusion

Improving the acquisition, transportation and utilization efficiency of Pi from soil to the plant is important for sustainable agriculture. Thus, it is essential to understand the mechanisms by which plants react and adapt to the P-deficiency. Our study identified important Pi responsive genes in maize by *in silico* analysis and characterized expression of these genes under Pi deficient conditions in a Pi stress tolerant genotype. Further studies may utilize the expression profile reported here, and investigate if over-expression of identified Pi-responsive genes can improve the PUE in maize.

## Author contributions

PY and TK conceived the idea and provided overall supervision to the study. VD and AA performed the experiments and analyzed the data with guidance from PY, IS and KK. RV helped establish the hydroponics. VD and AA wrote the primary draft, which was further augmented, edited and improved by PY. All the authors read and approved it for publication.

## Acknowledgements

This work was funded by a National Agricultural Science Fund grant (NASF/GTR-5004/2015-16/204) to PY and TK. The funders had no role in study design, data collection and analysis, decision to publish, or preparation of the manuscript.

## SUPPLEMENTARY MATERIAL

**Supplementary Table S1** Primers used for the expression analyses of 12 phosphate responsive genes of *Zea mays* L.

**Supplementary Table S2:** Identified phosphate starvation responsive gene homologs from *Zea mays* based on sequence similarity with *Arabidopsis* genes

**Supplementary Fig. S1** Domain analysis of putative PHO pathway regulatory genes in maize *viz*. PHR1 (phosphate starvation response 1), SPX1 domain containing protein, SIZ1, Pho2, ZAT6.

**Supplementary Fig. S2** Domain analysis of putative purple acid phosphatases (PAPs) and ribonuclease (RNase) in maize.

**Supplementary Fig. S3** Domain analysis of putative phosphate transporters in maize.

**Supplementary Fig. S4** Conserved PAP motifs in selected maize purple acid phosphatases (PAPs). Five conserved motifs, including DXG, GDXXY, GNH(D/E), VXXH, and GHXH are shown in maize and corresponding *Arabidopsis* PAPs using cobalt Multiple Alignment Tool from NCBI.

## References

Ai P, Sun S, Zhao J, Fan X, Xin W, Guo Q, Yu L, Shen Q, Wu P, Miller A (2009) Two rice phosphate transporters, OsPht1; 2 and OsPht1; 6, have different functions and kinetic properties in uptake and translocation. Plant J 57:798–809. https://doi.org/10.1111/j.1365-313X.2008.03726.x

Aung K, Lin SI, Wu CC, Huang YT, Su CL, Chiou TJ (2006) pho2, a phosphate overaccumulator, is caused by a nonsense mutation in a microRNA399 target gene. Plant Physiol 141:1000–1011. https://doi.org/10.1104/pp.106.078063

Bari R, Datt Pant B, Stitt M, Scheible WR (2006) PHO2, microRNA399, and PHR1 define a phosphate-signaling pathway in plants. Plant Physiol 141:988–999. https://doi.org/10.1104/pp.106.079707

Bariola PA, Howard CJ, Taylor CB, Verburg MT, Jaglan VD, Green PJ (1994) The Arabidopsis ribonuclease gene RNS1 is tightly controlled in response to phosphate limitation. Plant J 6:673–685. https://doi.org/10.1046/j.1365-313X.1994.6050673.x

Borevitz JO, Xia Y, Blount J, Dixon RA, Lamb C (2000) Activation tagging identifies a conserved MYB regulator of phenylpropanoid biosynthesis. Plant Cell 12:2383–2394. https://doi.org/10.1105/tpc.12.12.2383

Bozzo GG, Dunn EL, Plaxton WC (2006) Differential synthesis of phosphate-starvation inducible purple acid phosphatase isozymes in tomato (Lycopersiconesculentum) suspension cells and seedlings. Plant Cell and Env 29:303–313. https://doi.org/10.1111/j.1365-3040.2005.01422.x

Bindraban PS, Dimkpa CO, Pandey R (2020) Exploring phosphorus fertilizers and fertilization strategies for improved human and environmental health. Biology and Fertility of Soils 8:1–9.

Bustos R, Castrillo G, Linhares F, Puga MI, Rubio V, Pérez-Pérez J, Solano R, Leyva A, Paz-Ares J (2010) A central regulatory system largely controls transcriptional activation and repression responses to phosphate starvation in Arabidopsis. PLoS Genet 6:e1001102. https://doi.org/10.1371/journal.pgen.1001102

Chen YF, Li LQ, Xu Q, Kong YH, Wang H, Wu WH (2009) The WRKY6 transcription factor modulates PHOSPHATE1 expression in response to low Pi stress in Arabidopsis. Plant Cell 21:3554–3566. https://doi.org/10.1105/tpc.108.064980

Chen ZH, Nimmo GA, Jenkins GI, Nimmo HG (2007) BHLH32 modulates several biochemical and morphological processes that respond to Pi starvation in Arabidopsis. Biochem J 405:191–198. https://doi.org/10.1042/BJ20070102

Chiou TJ, Aung K, Lin SI, Wu CC, Chiang SF, Su CL (2006) Regulation of phosphate homeostasis by MicroRNA in Arabidopsis. Plant Cell 18:412–421. https://doi.org/10.1105/tpc.105.038943

Chowdhury RB, Zhang X (2021) Phosphorus use efficiency in agricultural systems: A comprehensive assessment through the review of national scale substance flow analyses. Ecological Indicators: 121:107172.

Cordell D, Drangert JO, White S (2009) The story of phosphorus: global food security and food for thought. Glob Environ Change 19(2):292–305. https://doi.org/10.1016/j.gloenvcha.2008.10.009

Deshpande RA, Shankar V (2002) Ribonucleases from T2 family. Crit Rev Microbiol 28:79–122. https://doi.org/10.1080/1040-840291046704

Devaiah BN, Madhuvanthi R, Karthikeyan AS, Raghothama, KG (2009) Phosphate starvation responses and gibberellic acid biosynthesis are regulated by the MYB62 transcription factor in Arabidopsis. Mol Plant 2:43–58. https://doi.org/10.1093/mp/ssn081

Devaiah BN, Nagarajan VK, Raghothama KG(2007) Phosphate homeostasis and root development in Arabidopsis are synchronized by the zinc finger transcription factor ZAT6. Plant Physiol 145(1):147–159. https://doi.org/10.1104/pp.107.101691

Dodds PN, Clarke AE,Newbigin E (1996) Molecular characterisation of an S-like RNase of Nicotianaalata that is induced by phosphate starvation. Plant Mol Biol 31: 227–238. https://doi.org/10.1007/BF00021786

Duff SMG, Sarath G, Plaxton WC (1994) The role of acid phosphatase in plant phosphorus metabolism. Plant Physiol 90:791–800. https://doi.org/10.1111/j.1399-3054.1994.tb02539.x

Feng S, Lim BL (2011) Functional role of a purple acid phosphatase in Arabidopsis thaliana. Degree of Doctor of Philosophy at The University of Hong Kong, pp 23–63.

Finn RD,Mistry J, Schuster-Böckler B, Griffiths-Jones S, Hollich V, Lassmann T et al. (2006) Pfam: clans, web tools and services. Nucleic Acids Res 34:D247–D251. https://doi.org/10.1093/nar/gkj149

Flanagan JU,Cassady AI, Schenk G, Guddat LW, Hume DA (2006) Identification and molecular modeling of a novel, plant-like, human purple acid phosphatase. Gene 377:12–20. https://doi.org/10.1016/j.gene.2006.02.031

Franco-Zorrilla JM, Gonzalez E, BustosR, LinharesF, Leyva A, Paz-Ares J (2004) The transcriptional control of plant responses to phosphate limitation. J Exp Bot 55:285–293. https://doi.org/10.1093/jxb/erh009

Fujii H, Chiou TJ, Lin SI, Aung K, Zhu JK (2005) AmiRNA involved in phosphate-starvation response in Arabidopsis. Curr Biol 15:2038–2043. https://doi.org/10.1016/j.cub.2005.10.016

Furihata T, Suzuki M, Sakurai H (1992) Kinetic characterization of two phosphate uptake systems with different affinities in suspension-cultured Catharanthus roseus protoplasts. Plant Cell Physiol 33:1151–1157. https://doi.org/10.1093/oxfordjournals.pcp.a078367

Ganie AH, Ahmad A, Pandey R, et al. (2015) Metabolite profiling of low-P tolerant and low-P sensitive maize genotypes under phosphorus starvation and restoration conditions. Plos one 10(6):e0129520. doi:10.1371/journal.pone.0129520.

González-Muñoz E, Avendaño-Vázquez AO, Montes RAC, De-Folter S, Andrés-Hernández L, Abreu-Goodger C, Sawers RJH (2015) The maize (Zea mays ssp. mays var. B73) genome encodes 33 members of the purple acid phosphatase family. Front Plant Sci 6:341. https://doi.org/10.3389/fpls.2015.00341

Gu M, Chen A, Sun S, Xu G (2016) Complex Regulation of plant phosphate transporters and the gap between molecular mechanisms and practical application: What is missing? Mol Plant 9:396–416. https://doi.org/10.1016/j.molp.2015.12.012

Higo K, Ugawa Y, Iwamoto M, Korenaga T (1999) Plant cis-acting regulatory DNA elements (PLACE) database: 1999. Nucleic Acids Res 27:297–300. https://doi.org/10.1093/nar/27.1.297

Huang CY, RoessnerU, EickmeierI, Genc Y, Callahan DL, Shirley N, Langridge P, Bacic A (2008) Metabolite profiling reveals distinct changes in carbon and nitrogen metabolism in phosphate-deficient barley plants (Hordeum vulgare L.). Plant Cell Physiol 49:691–703. https://doi.org/10.1093/pcp/pcn044

Jia H, Ren H, Gu M, ZhaoJ, SunS, Zhang X, ChenJ, WuP, Xu G (2011) The phosphate transporter gene OsPht1; 8 is involved in phosphate homeostasis in rice. Plant Physiol 156: 1164–1175. https://doi.org/10.1104/pp.111.175240

Jiang H, Zhang J, Han Z, Yang J, Ge C, Wu Q (2017) Revealing new insights into different phosphorus-starving responses between two maize (Zea mays) inbred lines by transcriptomic and proteomic studies. Sci Rep 7:44294. doi: 10.1038/srep44294.

Kock M, Loffler A, Abel S, Glund K (1995) cDNA structure and regulatory properties of a family of starvation-induced ribonucleases from tomato. Plant Mol Biol 27:477–485.

KockM, Stenzel I, Zimmer A (2006) Tissue-specific expression of tomato Ribonuclease LX during phosphate starvation-induced root growth. J Exp Bot 57:3717–3726. https://doi.org/10.1093/jxb/erl124

Kurepa J, Walker JM, Smalle J, Gosink MM, Davis SJ, Durham TL, Sung DY, Vierstra RD (2003) The small ubiquitin-like modifier (SUMO) protein modification system in Arabidopsis. Accumulation of SUMO1 and −2 conjugates is increased by stress. J Biol Chem 278:6862–6872. DOI: 10.1074/jbc.M209694200

Lescot M, Déhais P, Moreau Y, De Moor B, Rouzé P, Rombauts S (2002) PlantCARE: a database of plant cis-acting regulatory elements and a portal to tools for in silico analysis of promoter sequences. Nucleic Acids Res Database issue 30(1):325–327. https://doi.org/10.1093/nar/30.1.325

Letunic I, Doerks T, Bork P (2009) SMART 6: precent updates and new developments. Nucleic Acids Res 37:D229–D232. https://doi.org/10.1093/nar/gkn808

Li Z, Gao Q,Liu Y, He C, Zhang X, Zhang J (2011) Overexpression of transcription factor ZmPTF1 improves low phosphate tolerance of maize by regulating carbon metabolism and root growth. Planta 233:1129–1143. http://dx.doi.org/10.1007/s00425-011-1404-1

Li M, Tadano T (1996) Comparison of characteristics of acid phosphatases secreted from roots of lupin and tomato. Soil Sci Plant Nutr 42:753–763. https://doi.org/10.1080/00380768.1996.10416623

Li K, Xu Z, Zhang K, Yang A, Zhang J (2007) Efficient production and characterization for maize inbred lines with low-phosphorus tolerance. Plant Sci 172:255e264. https://doi.org/10.1016/j.plantsci.2006.09.004

Liang C, Tian J, Lam HM, LimBL, YanX, Liao H (2010) Biochemical and molecular characterization of PvPAP3, a novel purple acid phosphatase isolated from common bean enhancing extracellular ATP utilization. Plant Physiol 152:854–865. https://doi.org/10.1104/pp.109.147918

Liu C, Muchhal US, Uthappa M, Kononowicz AK, Raghothama KG (1998) Tomato phosphate transporter genes are differentially regulated in plant tissues by phosphorus. Plant Physiol 116: 91–99. https://doi.org/10.1104/pp.116.1.91

Liu F, Wang Z, Ren H, Shen C, Li Y, Ling HQ, Wu C, Lian X, Wu P (2010) OsSPX1 suppresses the function of OsPHR2 in the regulation of expression of OsPT2 and phosphate homeostasis in shoots of rice. Plant J 62:508–517. https://doi.org/10.1111/j.1365-313X.2010.04170.x

Liu PD, Xue Y B, Chen ZJ, Liu G D, Tian J (2016) Characterization of purple acid phosphatases involved in extracellular dNTP utilization in Stylosanthes. J Exp Bot 67:4141–4154. https://doi.org/10.1093/jxb/erw190

Liu P, Chen S, Song A, Zhao S, Fang W, Guan Z, Chen F (2014) A putative high affinity phosphate transporter, CmPT1, enhances tolerance to Pi deficiency of chrysanthemum. BMC plant biology 14:18. https://doi.org/10.1186/1471-2229-14-18

Lopez-Arredondo DL, Leyva-González MA, González-Morales SI, López-Bucio J, Herrera-Estrella L (2014) Phosphate nutrition: improving low-phosphate tolerance in crops. Annu Rev Plant Biol 65:95–123. https://doi.org/10.1146/annurev-arplant-050213-035949

Lun SC, Leung A, Kuang R, Wang Y, Leung P, Lim BL (2008) Phytase activity in tobacco (Nicotiana tabacum) root exudates is exhibited by a purple acid phosphatase. Phytochemistry 69:365–373. https://doi.org/10.1016/j.phytochem.2007.06.036

MacIntosh GC, Hillwig MS, Meyer A, Flagel L(2010) RNase T2 genes from rice and the evolution of secretory ribonucleases in plants. Mol Genet Genomics 283:381–396. https://doi.org/10.1007/s00438-010-0524-9

Matange N, Podobnik M, Visweswariah S (2015) Metallophosphoesterases: Structural fidelity with functional promiscuity. Biochem J 467:201–16. https://doi.org/10.1042/BJ20150028

Miller SS, Liu J, Allan DL, Menzhuber CJ,Fedorova M, Vance CP (2001) Molecular control of acid phosphatase secretion into the rhizosphere of proteoid roots from phosphorus-stressed white lupin. Plant Physiol 127:594–606. https://doi.org/10.1104/pp.010097

Misson J, Raghothama KG, Jain A, Jouhet J, Block MA, Bligny R, Ortet P, Creff A, Somerville S, Rolland N et al. (2005) A genome-wide transcriptional analysis using Arabidopsis thaliana Affymetrix gene chips determined plant responses to phosphate deprivation. Proc Natl Acad Sci USA 102:11934–11939. https://doi.org/10.1073/pnas.0505266102

Miura K, Jin JB, Lee J, Yoo CY, Stirm V, Miura T, Ashworth EN, Bressan RA, Yun DJ, Hasegawa PM (2007) SIZ1-mediated sumoylation of ICE1 controls CBF3/DREB1A expression and freezing tolerance in Arabidopsis. Plant Cell 19:1403–1414. https://doi.org/10.1105/tpc.106.048397

Miura K, Rus A, Sharkhuu A, Yokoi S, Karthikeyan AS, Raghothama KG, Baek D, Koo YD, Jin JB,Bressan RA, Yun DJ, Hasegawa PM (2005) The Arabidopsis SUMO E3 ligase SIZ1 controls phosphate deficiency responses. Proc Natl Acad Sci USA 102:7760–7765. https://doi.org/10.1073/pnas.0500778102

Müller R, Morant M, Jarmer H, Nilsson L, Nielsen TH (2007) Genome-wide analysis of the Arabidopsis leaf transcriptome reveals interaction of phosphate and sugar metabolism. Plant Physiol 143:156–171. https://doi.org/10.1104/pp.106.090167

Murphy J, Riley JP (1962) A modified single solution method for the determination of phosphate in natural waters. Anal Chim Acta 27:31–36. https://doi.org/10.1016/S0003-2670(00)88444-5

Nguyen GN, Rothstein SJ, Spangenberg G, Kant S (2015) Role of microRNAs involved in plant response to nitrogen and phosphorous limiting conditions. Front Plant Sci 6:629. https://doi.org/10.3389/fpls.2015.00629

Nilsson L, Müller R, Nielsen TH (2010) Dissecting the plant transcriptome and the regulatory responses to phosphate deprivation. Physiologia Plantarum 139:129–14310. https://doi.org/10.1111/j.1399-3054.2010.01356.x

NussaumeL, KannoS, Javot H, Marin E, PochonN, AyadiA, Nakanishi TM, ThibaudM C (2011) Phosphate import in plants: focus on the PHT1transporters. Front Plant Sci 2:83. https://doi.org/10.3389/fpls.2011.00083

Oropeza-Aburto A, Cruz-Ramírez A, Acevedo-Hernández GJ, Pérez-Torres CA, Caballero-Pérez J, Herrera-Estrella L (2012) Functional analysis of the Arabidopsis PLDZ2 promoter reveals an evolutionarily conserved low-Pi-responsive transcriptional enhancer element. J Exp Bot 63:2189–220210. https://doi.org/10.1093/jxb/err446

Ozawa K, Osaki M, Matsui H, Honma M, Tadano T (1995) Purification and properties of acid phosphatase secreted from lupin roots under phosphorus-deficiency conditions. Soil Sci Plant Nutr 41:461–469. https://doi.org/10.1080/00380768.1995.10419608

Peret B, Clement M, Nussaume L, Desnos T (2011) Root developmental adaptation to phosphate starvation: better safe than sorry. Trends Plant Sci 16:442–450. https://doi.org/10.1016/j.tplants.2011.05.006

Pozo DJC, AllonaI, RubioV, Leyva A, DeLa Peña A, Aragoncillo C, Paz-Ares J (1999) A type 5 acid phosphatase gene from Arabidopsis thaliana is induced by phosphate starvation and by some other types of phosphate mobilising/oxidative stress conditions. Plant J 19(5):579–589. https://doi.org/10.1046/j.1365-313X.1999.00562.x

Puga MI, Mateos I, Charukesi R, Wang Z, Franco-Zorrilla JM, de Lorenzo L, Leyva A (2014) SPX1 is a phosphate-dependent inhibitor of PHOSPHATE STARVATION RESPONSE 1 in Arabidopsis. Proc Natl Acad Sci 111:14947–14952. https://doi.org/10.1073/pnas.1404654111

Rae AL, Cybinski DH, Jarmey JM, Smith FW (2003) Characterization of two phosphate transporters from barley: evidence for diverse function and kinetic properties among members of the Pht1 family. Plant Mol Biol 53:27–36. https://doi.org/10.1023/B:PLAN.0000009259.75314.15

Raghothama K (2000) Phosphate transport and signaling. Curr Opin Plant Biol 3:182–187. DOI:10.1016/S1369-5266(00)00062-5

Raghothama KG (1999) Phosphate acquisition. Annu Rev Plant Physiol Plant MolBiol 50:665–693. https://doi.org/10.1146/annurev.arplant.50.1.665

Roalson EH, McCubbin AG (2003) S-RNases and sexual incompatibility: structure, functions, and evolutionary perspectives. Mol Phylogenet Evol 29:490–506. https://doi.org/10.1016/S1055-7903(03)00195-7

Robinson WD, Park J, Tran HT, Vecchio HAD, Ying S, Zins JL, Patel K,McKnight TD, Plaxton WC (2012) The secreted purple acid phosphatase isozymes AtPAP12 and AtPAP26 play a pivotal role in extracellular phosphate-scavenging by Arabidopsis thaliana. J Exp Bot 63:6531–42. https://doi.org/10.1093/jxb/ers309

Rouached H, Bulak A, Yves P (2010) Regulation of phosphate starvation responses in plants: signaling players and cross-talks. Mol Plant 3:288–299. https://doi.org/10.1093/mp/ssp120

Rubio V, Linhares F, Solano R, Martín AC, Iglesias J, LeyvaA, Paz-AresJ (2001) A conserved MYB transcription factor involved in phosphate starvation signaling both in vascular plants and in unicellular algae. Genes Dev 15:2122–2133. http://www.genesdev.org/cgi/doi/10.1101/gad.204401

Schenk G, Mitić N, Hanson GR, Comba P (2013) Purple acid phosphatase: a journey into the function and mechanism of a colorful enzyme. Coord Chem Rev 257:473–482. https://doi.org/10.1016/j.ccr.2012.03.020

Schnable PS, Ware D, Fulton RS, Stein JC, Wei F, Pasternak S, Liang C, Zhang J, Fulton L, Graves TA et al. (2009) The B73 maize genome: complexity, diversity, and dynamics. Sci 326:1112–1115. https://doi.org/10.1126/science.1178534

Schünmann PH, Richardson AE, Smith FW, Delhaize E (2004) Characterization of promoter expression patterns derived from the Pht1 phosphate transporter genes of barley (Hordeumvulgare L.). J Exp Bot 55:855–865. https://doi.org/10.1093/jxb/erh103

Shin H, Shin HS, Dewbre GR, Harrison MJ (2004) Phosphate transport in Arabidopsis: Pht1;1 and Pht1;4 play a major role in phosphate acquisition from both low- and high-phosphate environments. Plant J 39:629–642. https://doi.org/10.1111/j.1365-313X.2004.02161.x

Sun F, Suen PK, Zhang Y, Liang C, Carrie C, Whelan J, Lim BL(2012) A dual-targeted purple acid phosphatase in Arabidopsis thaliana moderates carbon metabolism and its overexpression leads to faster plant growth and higher seed yield. New Phytol 194:206–219. https://doi.org/10.1111/j.1469-8137.2011.04026.x

Taylor CB, Bariola PA, Delcardayre SB, Raines RT, Green PJ (1993) RNS2 - A senescence-associated RNase of Arabidopsis that divered from the S-RNases before speciation. Proc Natl Acad Sci USA 90:5118–5122. https://doi.org/10.1073/pnas.90.11.5118

Tian J, Liao H (2015) The role of intracellular and secreted purple acid phosphatases in plant phosphorus scavenging and recycling. Annu Plant Rev 48:265–287. https://doi.org/10.1093/jxb/ers309

Tohge T, Nishiyama Y, Hirai MY, Yano M, Nakajima J, Awazuhara M et al. (2005) Functional genomics by integrated analysis of metabolome and transcriptome of Arabidopsis plants over-expressing an MYB transcription factor. Plant J 42:218–35. https://doi.org/10.1111/j.1365-313X.2005.02371.x

Tran HT, Hurley BA, Plaxton WC (2010) Feeding hungry plants: the role of purple acid phosphatases in phosphate nutrition. Plant Sci 179:14–27. https://doi.org/10.1016/j.plantsci.2010.04.005

Udert KM (2018) Phosphorus as a resource. Phosphorus: Polluter and Resource, IWA Publishing, p. 57.

Veljanovski V, Vanderbeld B, Knowles VL, Snedden WA, Plaxton WC (2006) Biochemical and molecular characterization of AtPAP26, a vacuolar purple acid phosphatase up-regulated in phosphate-deprived Arabidopsis suspension cells and seedlings. Plant Physiol 142:1282–1293. https://doi.org/10.1104/pp.106.087171

Wang C, Ying S, Huang H, Li K, Wu P, Shou H (2009) Involvement of OsSPX1 in phosphate homeostasis in rice. Plant J 57:895–904. https://doi.org/10.1111/j.1365-313X.2008.03734.x

Wang L, Li Z, Qian W, Guo W, Gao X, Huang L, Liu D (2011) The Arabidopsis purple acid phosphatase AtPAP10 is predominantly associated with the root surface and plays an important role in plant tolerance to phosphate limitation. Plant Physiol 157(3):1283–1299. https://doi.org/10.1104/pp.111.183723

Wang Z, Ruan W, Shi J, Zhang L, Xiang D, Yang C et al. (2014) Rice SPX1 and SPX2 inhibit phosphate starvation responses through interacting withPHR2 in a phosphate-dependent manner. Proc Natl Acad Sci USA 111:14953–14958. https://doi.org/10.1073/pnas.1404680111

Wu Z, Zhao J, Gao R, Hu G, Gai J, Xu G, Xing H (2011) Molecular cloning, characterization and expression analysis of two members of the Pht1 family of phosphate transporters in Glycine max. PLoS One 6:e19752. https://doi.org/10.1371/journal.pone.0019752

Yao ZF, Liang CY, Zhang Q, Chen ZJ, Xiao BX, Tian J, Liao H (2014) SPX1 is an important component in the phosphorus signalling network of common bean regulating root growth and phosphorus homeostasis. J Exp Bot 65(12):3299–310. https://doi.org/10.1093/jxb/eru183

Yi K, Wu Z, Zhou J, Du L, Guo L, Wu Y et al. (2005) OsPTF1, a novel transcription factor involved in tolerance to phosphate starvation in rice. Plant Physiol 138:2087–2096. https://doi.org/10.1104/pp.105.063115

Yuan H, Liu D (2008) Signaling components involved in plant responses to phosphate starvation. J Integr Plant Biol 50:849–859. https://doi.org/10.1111/j.1744-7909.2008.00709.x

Zhang J, Zhou X, Xu Y, Yao M, Xie F, Gai J, Li Y, Yang S (2016) Soybean SPX1 is an important component of the response to phosphate deficiency for phosphorus homeostasis. Plant Sci 248:82–91. https://doi.org/10.1016/j.plantsci.2016.04.010

Zhou J, Jiao FC, Wu ZC, Li Y, Wang X, He X, Zhong W, Wu P(2008) OsPHR2 is involved in phosphate-starvation signaling and excessive phosphate accumulation in shoots of plants. Plant Physiol 146(4):1673–1686. https://doi.org/10.1104/pp.107.111443

Zhu H, Qian W, LuX, LiD, LiuX, LiuK, Wang D (2005) Expression patterns of purple acid phosphatase genes in Arabidopsis organs and functional analysis of AtPAP23 predominantly transcribed in flower. Plant Mol Biol 59:581–594. http://doi.org/10.1007/s11103-005-0183-0

